# Characterization of an *Osmr* Conditional Knockout Mouse Model

**DOI:** 10.1101/2023.10.27.564474

**Authors:** Logan S. Schwartz, Ruth L. Saxl, Tim Stearns, Jennifer J. Trowbridge

## Abstract

Oncostatin M (OSM) is a member of the interleukin-6 (IL-6) family of cytokines and has been found to have distinct anti-inflammatory and pro-inflammatory properties in various cellular and disease contexts. OSM signals through two receptor complexes, one of which includes OSMRβ. To investigate OSM-OSMRβ signaling in adult hematopoiesis, we utilized the readily available conditional *Osmr*^fl/fl^ mouse model B6;129-*Osmr*^*tm1*.*1Nat*^/J, which is poorly characterized in the literature. This model contains loxP sites flanking exon 2 of the *Osmr* gene. We crossed *Osmr*^fl/fl^ mice to interferon-inducible *Mx1*-Cre, which is robustly induced in adult hematopoietic cells. We observed complete recombination of the *Osmr*^fl^ allele and loss of exon 2 in hematopoietic (bone marrow) as well as non-hematopoietic (liver, lung, kidney) tissues. Using a TaqMan assay with probes downstream of exon 2, *Osmr* transcript was lower in the kidney but equivalent in bone marrow, lung, and liver from *Osmr*^fl/fl^ *Mx1*-Cre versus *Mx1*-Cre control mice, suggesting that transcript is being produced despite loss of this exon. Western blots show that liver cells from *Osmr*^fl/fl^ *Mx1*-Cre mice had complete loss of OSMR protein, while bone marrow, kidney, and lung cells had reduced OSMR protein at varying levels. RNA-seq analysis of a subpopulation of bone marrow cells (hematopoietic stem cells) finds that some OSM-stimulated genes, but not all, are suppressed in *Osmr*^fl/fl^ *Mx1*-Cre cells. Together, our data suggest that the B6;129-*Osmr*^*tm1*.*1Nat*^/J model should be utilized with caution as loss of *Osmr* exon 2 has variable and tissue-dependent impact on mRNA and protein expression.

## INTRODUCTION

OSM, belonging to the IL-6 family, plays an important role in development, malignancy, and homeostasis of various tissue types^1-5^. OSM is primarily produced by mature activated hematopoietic cells such as monocytes, T lymphocytes, neutrophils, macrophages, and dendritic cells^4,6,7^. OSM was first isolated based on its ability to inhibit the proliferation of melanoma tumor cells, while having no significant effect on normal human fibroblasts^6^. Due to this unique characteristic, it was given the name oncostatin M, where “onco” referred to cancer, “statin” to its ability to inhibit cell proliferation, and “M” to melanoma. Despite the initial identification of OSM as a tumor suppressor, a pro-inflammatory role of OSM has been reported in colon cancer, breast cancer, pancreatic cancer, myeloma, brain tumors, chronic lymphocytic leukemia, hepatoblastoma and COVID-19^1,3,8^. OSM, like IL-6, has been linked to tumor growth and is often found in high concentrations in the serum and tumor sites of patients and animal models of cancer^1,3,8^. Significantly increased *Osm* transcript production has also been associated with aging in both the human thymus and mouse hematopoietic stem cells (HSCs) compared to young tissues^9,10^. Human OSM interacts with two distinct receptor complexes termed type I and type II. The type I receptor complex contains leukemia inhibitory factor receptor alpha (LIFR) and interleukin 6 signal transducer (IL6ST; GP130), while the type II receptor complex contains oncostatin M receptor (OSMRβ) and IL6ST^4,11-13^. While recent studies have shown that OSM has a lower affinity for the type I receptor complex than the type II receptor complex^14,15^, much remains unknown about how OSM interacts with these complexes in different contexts. An additional complexity in studying OSM signaling is that while human OSM is known to signal through both the type I and type II receptor complexes, mouse OSM is reported to signal only through the type II receptor complex^16-18^.

To understand the role of OSM signaling in aging and disease in a physiologically relevant *in vivo* setting, mouse knockout models are essential. Two existing mouse models are commonly used to investigate loss of receptor complex II signaling through knockout of *Osmr*. The first, *Osmr*^tm1Mtan^, was reported by Tanaka et al. in 2003^19^. This model disrupts the first coding exon of *Osmr* by knock-in of a lacZ and neomycin expression cassette into the proximal region and is non-conditional in nature^19^. These *Osmr*^-/-^ mice were viable and fertile, born at expected Mendelian ratios, and had normal adult body weights^19^. Knockout of *Osmr* was confirmed by loss of mRNA in lung tissue^19^. This model exhibited hematopoiesis phenotypes showing reduced numbers of erythroid and megakaryocytic progenitors and their mature counterparts^19^. However, these were attributed to both cell-autonomous and cell non-autonomous effects of *Osmr*^-/-^ on hematopoietic progenitor cells and non-hematopoietic cells in the bone marrow microenvironment^19^, which are best resolved by the use of a conditional knockout model. The second existing model, B6;129-*Osmr*^*tm1*.*1Nat*^/J, is a conditional knockout model in which loxP sites flank the first coding exon of the *Osmr* gene.

Additionally, adjacent to the second loxP site is an Frt site remaining after excision of a PGK-neomycin selectable marker. This model was originally designed to study *Osmr* in the retina and is available in The Jackson Laboratory’s mouse repository; however, characterization of the model has not been published by the donating investigators. To investigate cell-autonomous effects of *Osmr* conditional knockout on hematopoiesis, we obtained the B6;129-*Osmr*^*tm1*.*1Nat*^/J (referred to as *Osmr*^fl/fl^) model and crossed these mice to an interferon-inducible *Mx1*-Cre allele^20^, which is commonly used in the study of adult hematopoiesis. We characterized *Osmr*^fl/fl^ *Mx1*-Cre mice using classical methods such as the development of a PCR reaction to evaluate recombination of the *Osmr*^fl^ allele, real-time PCR and RNA sequencing for *Osmr* mRNA transcript, and Western blotting to evaluate OSMRβ at the protein level. We investigated both hematopoietic tissue (bone marrow) as well as non-hematopoietic tissues (liver, lung, kidney) in which OSMRβ is also expressed. Our results demonstrate variable alterations in mRNA and protein expression in various tissues isolated from *Osmr*^fl/fl^ *Mx1*-Cre compared to control *Mx1*-Cre mice.

## RESULTS

### DNA Recombination in Bone Marrow Cells, Lung, Liver, and Kidney Cells of *Osmr*^fl/fl^ *Mx1*-Cre Mice

To generate an inducible *Osmr* knockout model, the B6;129-*Osmr*^*tm1*.*1Nat*^/J strain was crossed to *Mx1*-Cre^20^. *Mx1*-Cre is an interferon-inducible cre recombinase that is highly efficient in expression in progenitor and mature cells of the adult hematopoietic lineages. Thus, in the *Osmr*^fl/fl^ *Mx1*-Cre model, the *Osmr*^fl^ allele should remain unrecombined with wild-type expression at the transcript and protein levels prior to administration of the double-stranded RNA poly(I:C), which then induces an interferon response and expression of cre recombinase. After crossing B6;129-*Osmr*^*tm1*.*1Nat*^/J to *Mx1*-Cre to generate *Osmr*^fl/fl^ *Mx1*-Cre and control *Mx1*-Cre animals, we genotyped three control and three experimental animals alongside positive (parental *Osmr*^fl/fl^ and *Mx1*-Cre mice) and negative (wild-type C57BL/6J) controls. At the *Osmr* locus, genomic DNA from *Osmr*^fl/fl^ *Mx1*-Cre mice contained a loxP site (230bp) and genomic DNA from control *Mx1*-Cre mice did not contain a loxP site (170bp), recognized by primers flanking exon 2 (**Figure 1A**). Genomic DNA from all *Osmr*^fl/fl^ *Mx1*-Cre and control *Mx1*-Cre mice were also positive for the transgenic cre allele (100bp) (**Figure 1B**).

**Figure 1.**
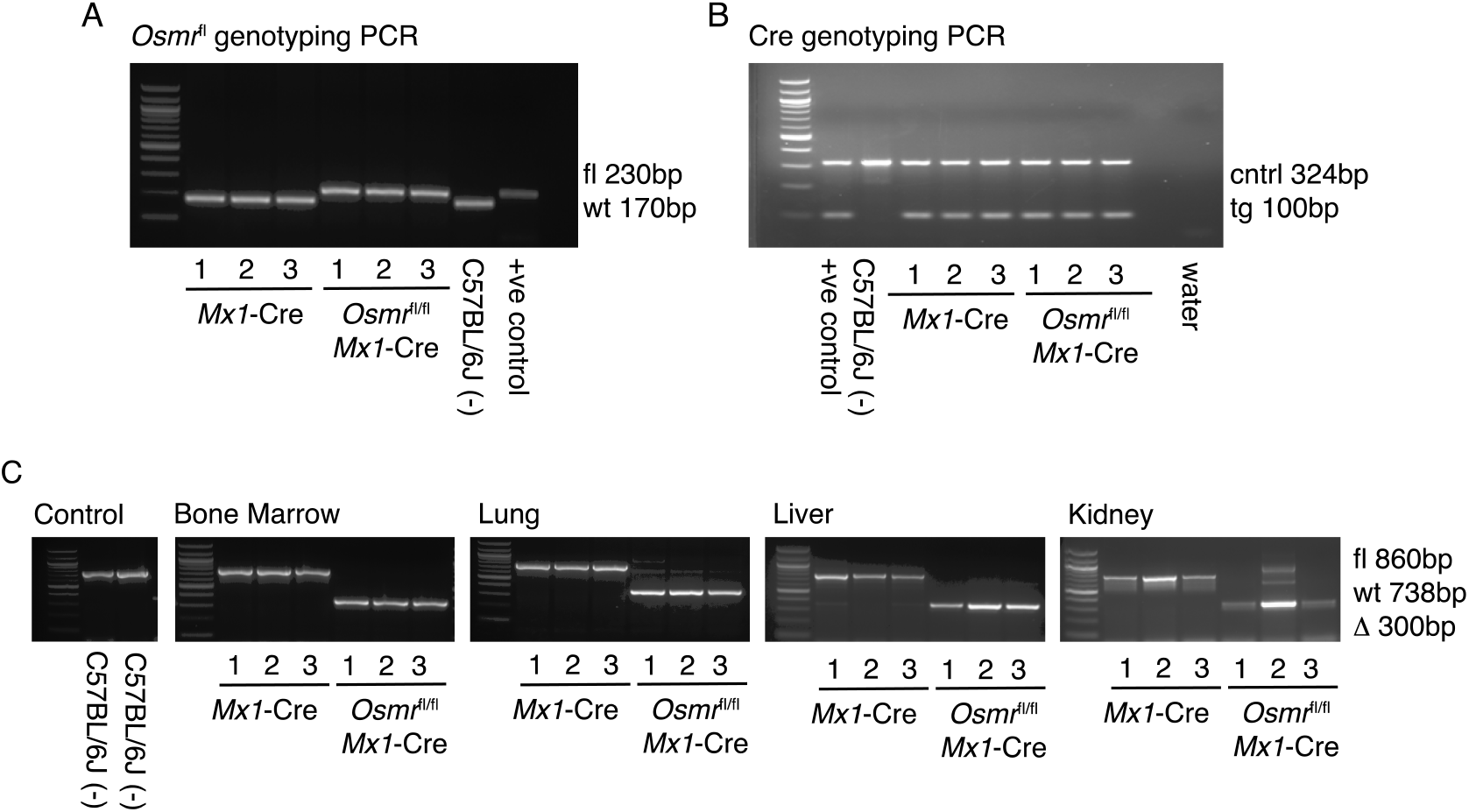
Recombination of the *Osmr*^fl^ Locus in Tissues from *Osmr*^fl/fl^ *Mx1*-Cre Mice. Agarose gel electrophoresis (2% agarose) of PCR-amplified products. In all gels a 100bp ladder was used for size estimation located in lane 0. **(A)** Genotyping PCR for Osmr<tm1.1Nat>. Lanes 1-3 use genomic DNA samples from *Mx1*-Cre control mice, lanes 4-6 use genomic DNA samples from *Osmr*^fl/fl^ *Mx1*-Cre mice, lane 7 uses genomic DNA from a wild-type C57BL/6J mouse and lane 8 uses a positive control *Osmr*^fl/fl^ from a mouse confirmed by The Jackson Laboratory’s in-house genotyping service. **(B)** Genotyping PCR for *Mx1*-Cre. Lane 1 shows a positive control, lane 2 shows wild-type C57BL/6J, lanes 3-5 show *Mx1*-Cre control mice and lanes 6-8 show *Osmr*^fl/fl^ *Mx1*-Cre mice. **(C)** Recombination PCR assay results in control, bone marrow, lung, liver, and kidney tissues. From left: Lanes 1-2 show wild-type C57BL/6J control mice. In the remaining images, samples are ordered as lanes 1-3 being *Mx1*-Cre control samples and lanes 4-6 being *Osmr*^fl/fl^ *Mx1*-Cre samples.

Following injection with poly(I:C) to induce *Mx1*-Cre-mediated recombination of the *Osmr*^fl^ locus, we isolated DNA from bone marrow, lung, liver, and kidney tissues. These tissues were selected based on expression of both *Osmr* and *Mx1* transcripts using publicly available databases including the human protein^21-23^ atlas, Mouse Genome Informatics^24^ and The Bgee suite^24^. A recombination PCR assay was designed to assess *Osmr*^fl^ recombination efficiency. The primers generate a product spanning the loxP-flanked exon 2 and bind to both the wild-type and modified *Osmr* loci generating product sizes of ∼860bp for the floxed (fl) allele, 738bp for the wild-type (wt) allele and ∼300bp for the recombined (*Δ*) allele. This PCR assay detected a ∼300bp recombined allele in all tissue samples from *Osmr*^fl/fl^ *Mx1*-Cre mice (**Figure 1C**), demonstrating that recombination of the *Osmr*^fl^ locus occurred efficiently following poly(I:C) injection in the bone marrow, lung, liver, and kidney.

### Tissue-Dependent Reduction in *Osmr* Transcript and Protein Expression *Osmr*^fl/fl^ *Mx1*-Cre Mice

We next evaluated levels of *Osmr* transcript in cells from the lung, liver, kidney, and bone marrow of *Osmr*^fl/fl^*Mx1*-Cre and control *Mx1*-Cre animals. Following harvest, cells were flash-frozen and used for RNA extraction, cDNA synthesis and semi-quantitative real-time PCR. TaqMan probes annealing to a target amplicon spanning *Osmr* exons 17 and 18, downstream of exon 2 deletion, were used. In control *Mx1*-Cre mice, *Osmr* transcript was detected at a low level in lung cells and robustly detected in liver, kidney, and bone marrow (**Figure 2A**). In *Osmr*^fl/fl^ *Mx1*-Cre mice, *Osmr* transcript in lung and liver was detected at similar levels as in control *Mx1*-Cre mice. In the kidney, *Osmr* transcript was significantly reduced in *Osmr*^fl/fl^ *Mx1*-Cre mice compared to control *Mx1*-Cre. In the bone marrow, *Osmr* transcript trended toward a decrease in *Osmr*^fl/fl^*Mx1*-Cre mice compared to control *Mx1*-Cre.

**Figure 2.**
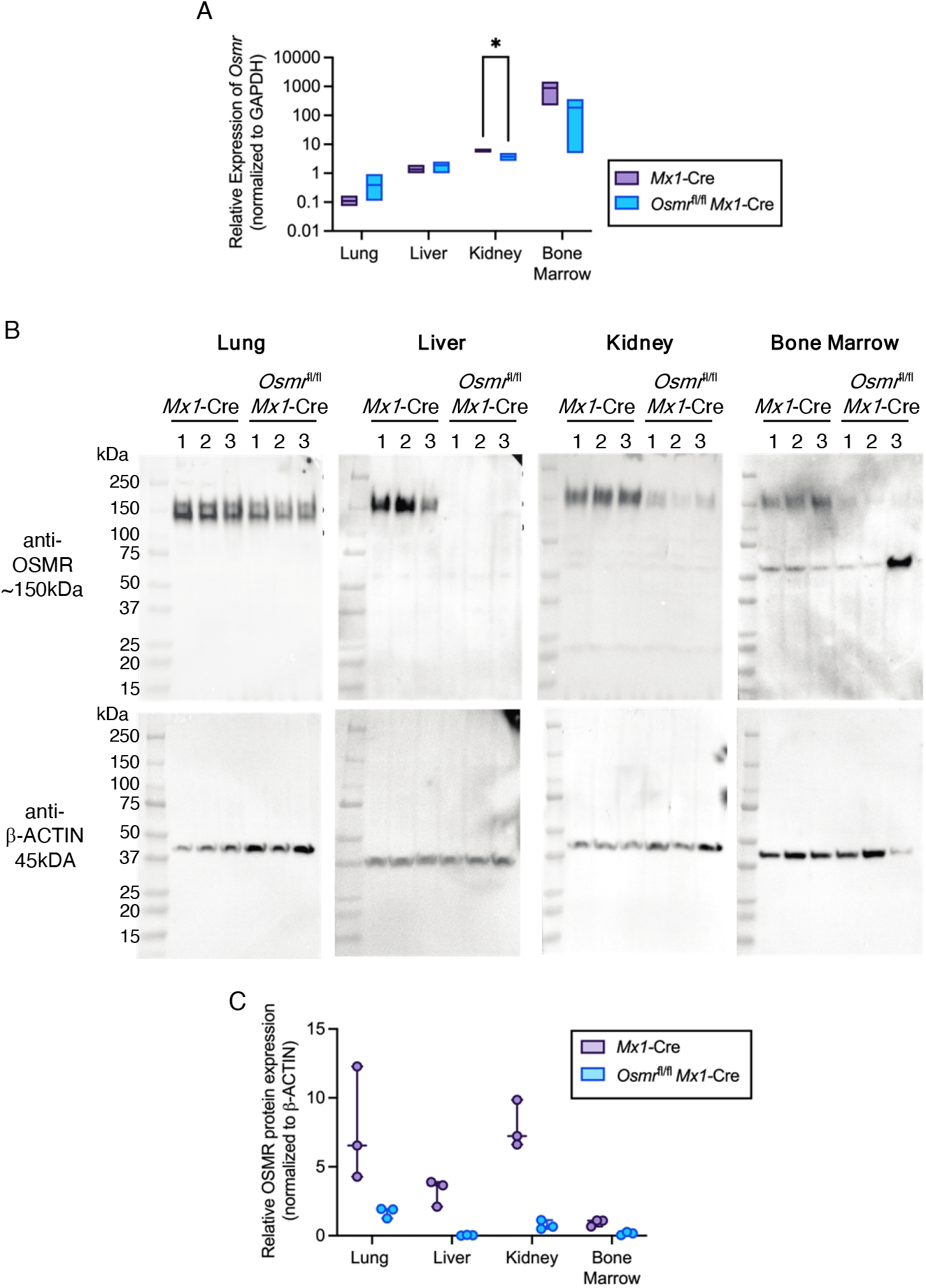
*Osmr*^fl/fl^ *Mx1*-Cre Mice Have Variable and Tissue-Dependent *Osmr* Expression at the Transcript and Protein Levels. **(A)** Relative *Osmr* expression assessed by real-time PCR in lung, liver, kidney, and bone marrow from *Mx1*-Cre control and *Osmr*^fl/fl^ *Mx1*-Cre mice (*n* = 3-4). Bars show low to high values with a line at the mean. \**p* < 0.05 by paired *t* test. **(B)** Western blots of cell lysates from lung, liver, kidney, and bone marrow probed with antibodies against OSMR and β-actin. **(C)** Relative protein expression assessed by Western blot imaging using ImageJ. OSMR band densities were normalized to β-actin (*n* = 3). Dots show individual bands/mice.

Given this tissue-dependent variation in *Osmr* transcript levels, we performed western blotting with a published OSMR antibody^14,16,25,26^ to quantify protein expression in the lung, liver, kidney, and bone marrow isolated from *Osmr*^fl/fl^ *Mx1*-Cre and control *Mx1*-Cre mice. OSMR protein expression showed modest reduction in the lung of *Osmr*^fl/fl^ *Mx1*-Cre mice compared to control *Mx1*-Cre mice (**Figure 2B-C**). OSMR expression in the liver was undetectable in *Osmr*^fl/fl^ *Mx1*-Cre mice, and OSMR expression in the kidney and bone marrow were reduced to very low levels in *Osmr*^fl/fl^ *Mx1*-Cre mice. Together, these data suggest that *Osmr*^fl/fl^ *Mx1*-Cre mice have reduced OSMR in cell lysates prepared from lung, liver, kidney, and bone marrow and that the level of reduction in OSMR protein is tissue dependent.

### Exploring Transcriptional Alterations in *Osmr*^fl/fl^ *Mx1*-Cre Hematopoietic Stem Cells (HSCs)

Given the reduction in OSMR protein detected in bone marrow lysates of *Osmr*^fl/fl^ *Mx1*-Cre mice, we utilized this model to investigate *Osmr* signaling in hematopoietic stem cells (HSCs) which are a small subset of bone marrow hematopoietic cells. HSCs were prospectively isolated from the bone marrow of mice using fluorescence-activated cell sorting. HSCs from *Osmr*^fl/fl^ *Mx1*-Cre and control *Mx1*-Cre mice were placed into serum-free media with or without 500ng/mL of recombinant murine OSM or vehicle control for 60min, then flash frozen for RNA sequencing (RNA-seq). Significantly differentially expressed genes were defined based on stringent cutoffs (*p* < 0.01 and log2(FC) > 3 or < -3) (**Figure 3A, Supplemental Table 1**). Consistent with our real-time PCR analysis of bone marrow cells, no difference in *Osmr* transcript abundance was observed in *Osmr*^fl/fl^ *Mx1*-Cre HSCs compared to control *Mx1*-Cre HSCs (**Figure 3B**). We removed predicted genes and pseudogenes from this list, resulting in a total of 91 significantly differentially expressed genes between *Osmr*^fl/fl^ *Mx1*-Cre and control *Mx1*-Cre HSCs. Of these, 59 genes were increased in expression and 32 were decreased in expression in *Osmr*^fl/fl^ *Mx1*-Cre HSCs. The top differentially expressed genes based on fold change included increased expression of *Gbp2b, Myh3, Cplx3, Gpr31c* and decreased expression of *Wdpcp, Lgalsl, Tesk2, Tigar*. Functional annotation of differentially expressed genes in *Osmr*^fl/fl^ *Mx1*-Cre compared to control *Mx1*-Cre HSCs identified enrichment of signatures related to proliferation, actin binding, RNA Pol II activity and protein binding, and depletion of signatures related to smoothened pathway signaling, intraciliary transport, and protein phosphorylation (**Figure 3C**). These data suggest that modest transcriptional differences are observed in HSCs isolated from *Osmr*^fl/fl^ *Mx1*-Cre compared to *Mx1*-Cre mice, despite lack of detectable differences in *Osmr* transcript itself.

**Figure 3.**
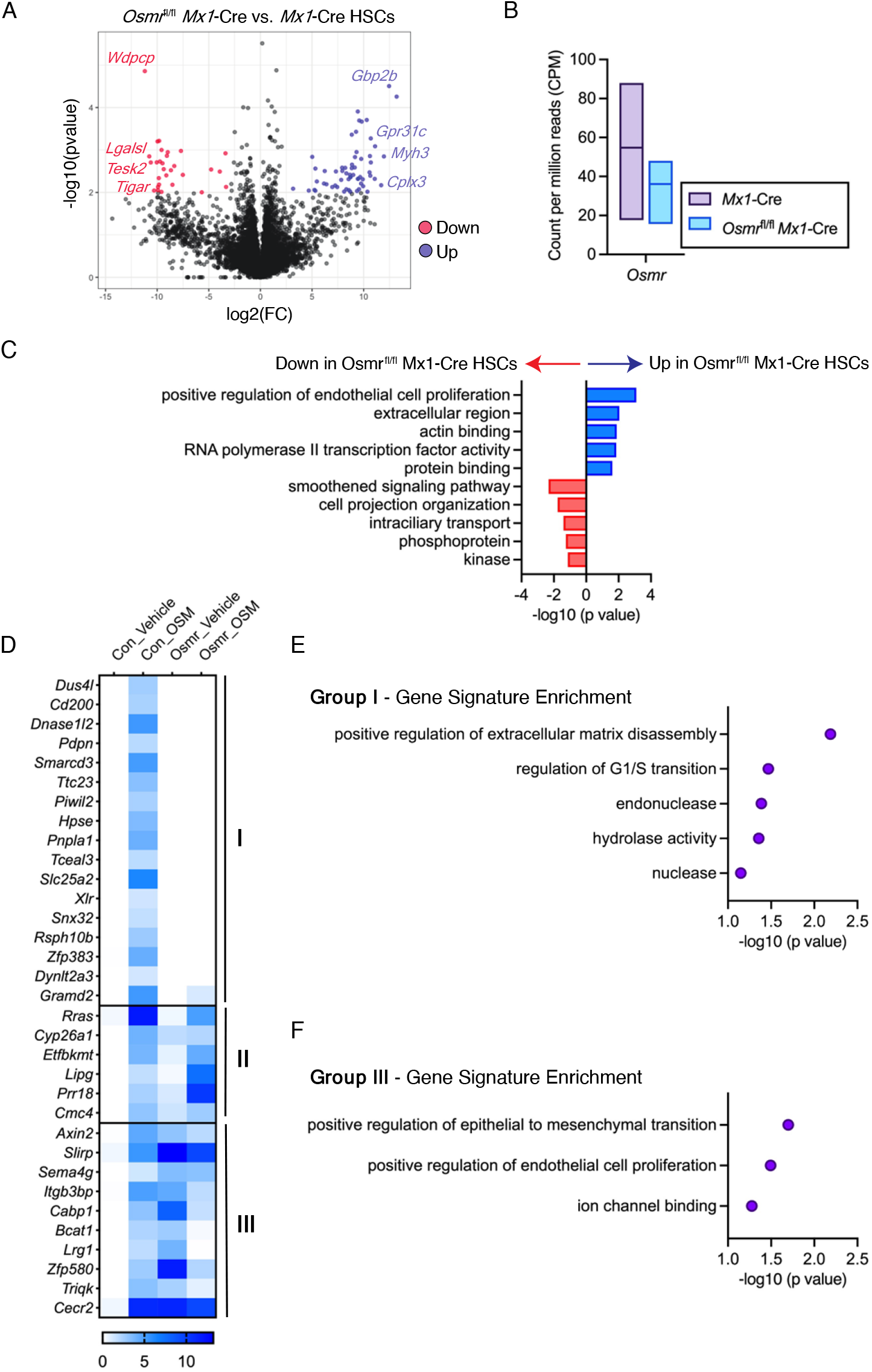
Differential Expression Analysis in *Osmr*^fl/fl^ *Mx1*-Cre HSCs. **(A)** Volcano plot showing differential gene expression of analysis of *Osmr*^fl/fl^ *Mx1*-Cre vs. control *Mx1*-Cre HSCs (*n* = 3-6 biological replicates). Select significantly differentially expressed genes (*p* < 0.01, log2(FC) > 3 or < -3) are shown in red (down) or blue (up) font. **(B)** Count per million reads of *Osmr* transcript in control *Mx1*-Cre and *Osmr*^fl/fl^ *Mx1*-Cre HSCs. Bars show low to high values with a line at the mean. **(C)** Gene signature enrichment within differentially expressed genes between *Osmr*^fl/fl^ *Mx1*-Cre and control *Mx1*-Cre HSCs. **(D)** Heatmap of 37 genes induced by 60min OSM stimulation in control *Mx1*-Cre HSCs. Three patterns were observed: Group (I) genes were no longer induced by OSM in *Osmr*^fl/fl^ *Mx1*-Cre HSCs, group (II) genes induced by OSM in both control *Mx1*-Cre and *Osmr*^fl/fl^ *Mx1*-Cre HSCs, and group (III) genes induced by OSM in control *Mx1*-Cre HSCs but repressed by OSM in *Osmr*^fl/fl^ *Mx1*-Cre HSCs. **(E)** Gene signature enrichment within group (I) genes. **(F)** Gene signature enrichment within group (III) genes.

Next, we utilized our RNA-seq dataset to explore how control *Mx1*-Cre versus *Osmr*^fl/fl^ *Mx1*-Cre HSCs respond to stimulation with recombinant OSM. Using the same stringent cutoffs defined above to identify significantly upregulated genes, control *Mx1*-Cre HSCs had 37 genes increased in expression following OSM stimulation compared to vehicle treatment (**Figure 3D, Supplemental Table 2**). Examining these 37 genes in *Osmr*^fl/fl^ *Mx1*-Cre HSCs stimulated with OSM (**Supplemental Table 3**) revealed several patterns; (I) 17 genes were no longer induced by OSM in *Osmr*^fl/fl^ *Mx1*-Cre HSCs, (II) 6 genes were induced by OSM in both control *Mx1*-Cre and *Osmr*^fl/fl^ *Mx1*-Cre HSCs, and (III) 10 genes were induced by OSM in control *Mx1*-Cre HSCs but repressed by OSM in *Osmr*^fl/fl^ *Mx1*-Cre HSCs. The 17 genes that were induced by OSM in control *Mx1*-Cre

HSCs but not induced in *Osmr*^fl/fl^ *Mx1*-Cre HSCs were enriched for signatures of extracellular matrix disassembly, G1/S transition, endonuclease, and hydrolase activity (**Figure 3E**). In contrast, the 10 genes that were induced by OSM in control *Mx1*-Cre HSCs but repressed by OSM in *Osmr*^fl/fl^ *Mx1*-Cre HSCs were enriched for signatures of epithelial to mesenchymal transition, endothelial cell proliferation and ion channel binding (**Figure 3F**). These data suggest that a subset of genes transcriptionally upregulated in response to OSM signaling in control *Mx1*-Cre HSCs are no longer upregulated in response to OSM in *Osmr*^fl/fl^ *Mx1*-Cre HSCs, consistent with genetic knockout and functional loss of OSMR. However, as a subset of OSM signaling target genes are increased in expression in *Osmr*^fl/fl^ *Mx1*-Cre HSCs, and other genes are no longer induced by OSM signaling, these results suggest that there is an incomplete loss of OSMRβ and/or that these genes are not *bona fide* targets of OSM-OSMR*β* signaling.

## DISCUSSION

Our data show that *Osmr*^fl/fl^ *Mx1*-Cre mice have near-complete recombination of the genomic *Osmr*^fl^ locus in bone marrow, lung, liver, and kidney tissue. However, the extent of reduction in *Osmr* transcript and OSMRβ protein expression in *Osmr*^fl/fl^ *Mx1*-Cre tissues compared to *Mx1*-Cre tissues was dependent upon the tissue type examined. These findings are consistent with a previous study of the B6;129-*Osmr*^*tm1*.*1Nat*^/J model showing reduced OSMRβ, as opposed to a complete knockout, in K5+ epidermis^14,16,25,26^. Additionally, despite the lack of observable differences in *Osmr* transcript expression in HSCs isolated from *Osmr*^fl/fl^ *Mx1*-Cre compared to *Mx1*-Cre mice, we observed alterations in the levels of other transcripts. Focusing on genes induced by OSM signaling in control *Mx1*-Cre HSCs, we found that only some of these genes are no longer induced by OSM signaling in *Osmr*^fl/fl^ *Mx1*-Cre HSCs. We interpret this data to mean that residual OSMRβ expression in *Osmr*^fl/fl^ *Mx1*-Cre HSCs permits some OSM signaling to occur. In addition, HSCs with reduction or loss of OSMRβ appear to incur compensatory activation of OSM-stimulated genes indicating a more complex, tissue-dependent regulation of OSM-OSMRβ signaling than was previously understood.

Future studies will be necessary to understand the mechanisms underlying tissue-dependent differences in *Osmr* transcript abundance and OSMRβ protein abundance in the B6;129-*Osmr*^*tm1*.*1Nat*^/J model. It is possible that tissue-dependent differences in transcript isoforms and/or splicing underly differences in abundance of transcript and protein. For example, transcripts lacking exon 2 may be sufficient to produce OSMRβ protein in the lung but not the liver or bone marrow. The appearance of two high molecular weight bands in the OSMRβ western blot in the lung but not in other tissues is consistent with multiple protein products being present in this tissue. In the bone marrow, a lower molecular weight band was visible in the OSMRβ western blot, which may indicate a shorter OSMRβ isoform that is still produced in *Osmr*^fl/fl^ *Mx1*-Cre cells.

Utilizing mouse models to understand the role of OSM signaling in human pathophysiology is essential but is an inherent challenge due to the previously characterized differences in receptor binding between mice and humans. Other studies have reported novel and innovative tools to aid in overcoming this challenge. For example, with the knowledge that specific charged residues in the AB loop of OSM are crucial for selection of binding to receptor complex I versus complex II, “humanized” murine OSM variants have been generated to gain more accurate insights into the role of human OSM in mouse models^16^. In addition, a potent and specific inhibitor of OSM signaling has been engineered using a receptor fusion protein that contains the ligand binding domains of murine OSMRβ and murine GP130^27^. This construct can be delivered into cells using viral vectors and has been used in multiple studies investigating the hematopoietic stem cell niche in the bone marrow and the role of OSM in irritable bowel disease^26,28^. Using this construct to generate transgenic animals would be very useful for future *in vivo* studies. Moving forward, we advise that other tools be utilized in parallel to the B6;129-*Osmr*^*tm1*.*1Nat*^/J model and/or that studies include a detailed characterization of *Osmr* transcript and protein levels in the tissue and cell type of focus.

## METHODS

### Animals

C57BL/6J (JAX:000664) mice were obtained from The Jackson Laboratory. B6;129-*Osmr*^*tm1*.*1Nat*^/J (JAX: 011081) were crossed to B6.Cg-Tg(*Mx1*-cre)1Cgn/J mice (JAX:003556). In all experiments, experimental and control mice carried a single copy of the *Mx1*-Cre transgenic allele. To induce *Mx1*-Cre, mice were intraperitoneally injected once every other day for five total injections with 15 mg/kg high molecular weight polyinosinic-polycytidylic acid (polyI:C) (InvivoGen). Mice used for experiments were at least 10-weeks following polyI:C administration. Tissues with medium-to-high expression of *Mx1* and *Osmr* were selected using publicly available databases such as the human protein atlas^21-23^, Mouse Genome Informatics^24^ and The Bgee suite^24^. The Jackson Laboratory’s Institutional Animal Care and Use Committee approved all experiments.

### Genotyping and Recombination PCR

Genomic DNA was extracted from lysed peripheral blood samples using the DNeasy Blood & Tissue Kit (Qiagen) for genotyping. Genotyping primers were used as suggested on the B6;129-*Osmr*^*tm1*.*1Nat*^/J (JAX: 011081) mice webpage (protocol ID 29086 and 23692). Images were taken using the InGenuis LHR2 Gel imaging system (Syngene). Recombination of the *Osmr*^fl^ locus was evaluated by PCR using the following primers: Forward: 5’-GGAAATACCTTGGCAGTGGTG and Reverse: 5’-GCTACCAAACCTCGGTAATCC.

### Osmr mRNA Expression

RNA was isolated using the RNeasy Minikit (Qiagen) and cDNA was made using the qPCR Bio cDNA synthesis kit (PCR Biosystems). Quantitative PCR was performed using TaqMan™ Fast Advanced Master Mix (Applied Biosystems) on the QuantStudio 7 Real-Time PCR System (ThermoFisher Scientific). mRNA expression levels for *Osmr* (Taqman probe #Mm01307326_m1) were calculated relative to the housekeeping gene, *Gapdh* (Taqman probe # Mm99999915_g1).

### RNA Sequencing

Bone marrow (BM) cells were isolated from *Osmr*^fl/fl^ *Mx1*-Cre and *Mx1*-Cre control mice from pooled and crushed femurs, tibiae, iliac crests, sternums, forepaws, and spinal columns of individual mice. BM mononuclear cells (MNCs) were isolated by 1X Red Blood Cell lysis Buffer (eBioscience) and stained with a combination of fluorochrome-conjugated antibodies: c-Kit (BD Biosciences, BioLegend clone 2B8), Sca-1 (BioLegend clone D7), CD150 (BioLegend clone TC15-12F12.2), CD48 (BioLegend clone HM48-1), FLT3 (BioLegend clone A2F10), (Lin) marker mix (B220 (BD Biosciences, BioLegend clone RA3-6B2), and the 4′,6-diamidino-2-phenylindole (DAPI). 2,000 hematopoietic stem cells (HSCs) were sorted based on the cell surface marker combination: Lin-Sca-1+ c-Kit+ Flt3-CD150+ CD48-on a FACSAria into StemSpan SFEM II media with SCF (100 ng/ml, BioLegend), TPO (50 ng/ml, Peprotech), with or without OSM (500 ng/ml, BioLegend) and incubated at 37°C for 60min. RNA preparation, sequencing and analysis was performed following a previously published method^29^. Significantly differentially expressed genes were identified based on criteria of *p* < 0.01 and log2(FC)>3 or <-3. Predicted genes and pseudogenes were removed from the list of significantly differentially expressed genes before downstream comparisons were performed. The DAVID bioinformatics resource^30,31^ was used for functional annotation of significantly differentially expressed gene lists.

### Western Blotting for OSMRβ

Total protein was extracted from snap frozen tissues with a Tissue Lyser II (Qiagen) using 5mm Bead Beater balls (Qiagen) after the addition of 50mM HEPES pH 7.5 (Gibco) and 6M urea (Sigma-Aldrich) at 1.5mL per 200mg tissue. Insoluble material was spun out at 21,000 x g for 20m. Total protein concentration of each sample was determined by microBCA (ThermoFisher). SDS-PAGE gel samples were prepared. Two 4–15% Mini-PROTEAN TGX Precast, 1.0mm, 15-well, SDS-PAGE Gel (Bio-Rad) were loaded with 25ug total protein per well. The gels were run at 150V for 1h at room temperature in Tris·Glycine·SDS buffer (Bio-Rad) then transferred overnight at 4°C at 40mAmp to 0.22mm PVDF (Bio-Rad) in Tris·Glycine buffer (Bio-Rad) with 10% methanol. The blots were placed in Block (2% BSA + 4% Dry Milk) at room temperature for 1h. The blots were then placed in primary antibody Polyclonal Goat IgG anti-mouse OSMR (R&D systems AF662) diluted in fresh block at 1:2500 and incubated on an orbital shaker at 4°C overnight. Then the blots were washed with TBS with 0.03% Tween 20 (TBS-T) for 15min and three more washes for 5 minutes each. Then the blots were placed in Peroxidase AffiniPure Bovine Anti-Goat IgG (H+L)(Jackson Immuno Research) diluted 1:5,000 in block. The blots rotated for 1h at room temperature and then were washed with TBS-T for 15min and three more washes for 5 minutes each.

The blot was then developed with (ThermoFisher) SuperSignal West Pico Plus reagents (ThermoFisher) and visualized with a Syngene G-Box. The blots were then stripped with Restore™ PLUS Western Blot Stripping Buffer (ThermoFisher) at room temperature for 5min. The blots were re-blocked and then incubated with β-actin (Cell Signaling Technology clone D6A8) at 1:5000 for 1h at room temperature. Then the blots were washed with TBS-T for 15min and three more washes for 5 minutes each. Then the blots were placed in Goat Anti-Rabbit IgG (H+L)-HRP (Bio-Rad) diluted 1:5,000 in block. The blots rotated for 1h at room temperature and then were washed with TBS-T for 15min and three more washes for 5 minutes each. The blot was then developed with (ThermoFisher) SuperSignal West Femto Maximum Sensitivity reagents (ThermoFisher) and visualized with a Syngene G-Box. Images were quantified by ImageJ by relative intensity, adjusted for background and normalized to β-ACTIN expression per sample.

## Data Availability

Raw RNA-seq data is available at the Gene Expression Omnibus under GSE236693 and GSE244544.

## AUTHORSHIP

L.S. and J.J.T. conceptualized the project and designed experiments. L.S. performed experiments, and L.S and J.J.T. analyzed data. R.S. performed Western blot experiments and image analysis. T.S. assisted in analysis of RNA-seq data. L.S. and J.J.T. wrote the manuscript.

## CONFLICT OF INTEREST DISCLOSURE

J.J.T. has previously received research support from H3 Biomedicine, Inc., and patent royalties from Fate Therapeutics.

## ACKNOWLEDGEMENTS

This work was supported by National Institutes of Health grants R01DK118072, R01AG069010, U01AG077925, and an EvansMDS Discovery Research Grant to J.J.T. This work was supported in part by the NIH/NCI Cancer Center Support Grant P30CA034196. J.J.T. was supported by a Leukemia & Lymphoma Society Scholar Award and The Dattels Family Endowed Chair. L.S.S. was supported by F31DK127573 and The Tufts University Scheer-Tomasso Fund philanthropic gift. We thank Drs. Jeremy Nathans and Amir Rattner for providing detailed information regarding the design of the B6;129-*Osmr*^*tm1*.*1Nat*^/J allele. We thank all members of the Trowbridge Lab, particularly Jayna Mistry, for experimental support and manuscript editing. We thank the Scientific Services at The Jackson Laboratory including flow cytometry and genome technologies. We thank Elli Hartig, Carol Bult, Ryan Tewhey, Cliff Rosen, and Phil Hinds for discussion and input into this work.

---

**Supplemental Table 1. Differential Gene Expression Analysis of *Osmr*^fl/fl^ *Mx1*-Cre vs. Control *Mx1*-Cre HSCs from RNA-Seq Data.** See Excel file.

**Supplemental Table 2. Differential Gene Expression Analysis of OSM-Stimulated vs. Vehicle-Treated Control *Mx1*-Cre HSCs from RNA-Seq Data.** See Excel file.

**Supplemental Table 3. Differential Gene Expression Analysis of OSM-Stimulated vs. Vehicle-Treated *Osmr*^fl/fl^ *Mx1*-Cre HSCs from RNA-Seq Data.** See Excel file.

